# Genomic balancing selection is key to the invasive success of the fall armyworm

**DOI:** 10.1101/2020.06.17.154880

**Authors:** Sudeeptha Yainna, Wee Tek Tay, Estelle Fiteni, Fabrice Legeai, Anne-Laure Clamens, Sylvie Gimenez, Marie Frayssinet, R Asokan, CM Kalleshwaraswamy, Sharanabasappa Deshmukh, Robert L. Meagher, Carlos A. Blanco, Pierre Silvie, Thierry Brévault, Anicet Dassou, Gael J. Kergoat, Thomas Walsh, Karl Gordon, Nicolas Nègre, Emmanuelle d’Alençon, Kiwoong Nam

## Abstract

A successful biological invasion involves survival in a newly occupied environment. If a population bottleneck occurs during an invasion, the resulting depletion of genetic variants could cause increased inbreeding depression and decreased adaptive potential, which may result in a fitness reduction. How invasive populations survive in the newly occupied environment despite reduced heterozygosity and how, in many cases, they maintain moderate levels of heterozygosity are still contentious issues^1^. The Fall armyworm (FAW; Lepidoptera: *Spodoptera frugiperda*), a polyphagous pest, is native to the Western hemisphere. Its invasion in the Old World was first reported from West Africa in early 2016, and in less than four years, it swept sub-Saharan Africa and Asia, finally reaching Australia. We used population genomics approaches to investigate the factors that may explain the invasive success of the FAW. Here we show that genomic balancing selection played a key role in invasive success by restoring heterozygosity before the global invasion. We observe a drastic loss of mitochondrial polymorphism in invasive populations, whereas nuclear heterozygosity exhibits a mild reduction. The population from Benin in West Africa has the lowest length of linkage disequilibrium amongst all invasive and native populations despite its reduced population size. This result indicates that balancing selection increased heterozygosity by facilitating the admixture of invasive populations from distinct origins and that, once heterozygosity was sufficiently high, FAW started spreading globally in the Old World. As comparable heterozygosity levels between invasive and native populations are commonly observed^1^, we postulate that the restoration of heterozygosity through balancing selection could be widespread among successful cases of biological invasions.

## Text

A successful biological invasion involves the survival of an introduced population, which is typically associated with rapid adaptation processes in the newly occupied environment^2,3^. If a bottleneck occurs during an invasion as a result of the introduction of a small number of individuals, the invasive population may have a decreased fitness due to inbreeding depression because the level of heterozygosity is decreased^4^. Moreover, small populations may have a lower adaptive potential than large populations because of a lower population-scaled rate of mutation^5–7^ or a lower number of existing genetic variants^8^, of which a proportion provides a beneficial effect for the survival in a new environment. The expectation that invasive populations have a reduced fitness appears to be contradictory with ample cases of invasive success, which has been often coined up as the ‘genetic paradox of biological invasion’^9^.

The occurrence of multiple introduction events has been proposed to be the solution to this paradox because genetic admixture among heterogeneous populations results in an increase in heterozygosity, which may decrease inbreeding depression and increase adaptive potential (reviewed in Estoup et al.^1^). However, the co-existence of allopatrically-originated individuals does not necessarily cause an increase in the level of heterozygosity because of the following two reasons. First, admixed individuals may have reduced fitness due to genetic incompatibilities between two haplotypes or strains. An established population is expected to have an optimal allelic combination through natural selection. Thus, admixed individuals between two established populations may have a substantial number of incompatible alleles, which decreases fitness. Indeed, genetic incompatibilities between populations are common in *Drosophila* fruit flies^10^. In addition, during the entire process of admixture, the stochastic effect of genetic drift may cause a substantial loss of variants if the initial number of invading individuals is small. In other words, a large effective population size is required to maintain variants from heterogeneous populations by overcoming genetic drift.

If selective advantages of admixed genotypes are sufficiently high to overcome the potential genetic incompatibilities in admixed individuals or to overcome genetic drift at the initial phase of admixture, then balancing selection may act in the way of facilitating admixture between different sets of genotypes from the different invasive origins. Therefore, it is tempting to hypothesize that invasive populations experience balancing selection, which restores heterozygosity during the lagging time between initial introductions to rapid range expansion.

The fall armyworm, *Spodoptera frugiperda* (J.E. Smith) (Lepidoptera: Noctuidae: Noctuinae), is one of the most infamous insect pests due to an extremely high-level of polyphagy (more than 353 host-plants belonging to 76 plant families are reported^11^), high dispersal capacity and migratory behavior^12^,the rapid development of insecticide resistance^13,14^, including resistance to Bt proteins^15–18^, and occasional outbreaks^19^. The FAW is native to North and South America, and its presence in West Africa was first reported in 2016^20^. In the following years, the FAW spread across sub-Saharan Africa, followed by global detection in India, South East Asia, East Asia, Egypt, and Australia (https://www.cabi.org/isc/fallarmyworm). Invasive FAW larvae cause significant economic losses, especially on corn, with yield loss of corn production averaging 21%-53% in Africa^21^. The FAW consists of two strains, corn strain (sfC) and rice strain (sfR) (named after their supposedly preferred host-plants), which are observed sympatrically in all their native range^22–24^. Both strains are observed in invasive populations, while the relative proportion of the identified strains depends on their geographic location^25–27^. Tay et al., reported genomic signature of multiple introductions of FAW from Mississippi and South America to the Old World based on 870 unlinked single nucleotide variants (SNV)^28^. Potential multiple introductions and the recent explosive global invasion of the FAW makes this species an ideal model to test the potential effect of balancing selection in invasion success.

In this paper, we aim at testing the potential role of balancing selection in the global invasion of the FAW using population genomics. First, we identified genomic SNV (Single Nucleotide Variants) from 177 samples in both native and invasive populations. Then, we inferred multiple origins of invasion, and tested balancing selection in the invasive population. Lastly, we identified adaptive evolution specific to invasive populations. We generated a new reference genome assembly from sfC using 30X PacBio Reads and Hi-C data^29^. The assembly size and N50 are 385 Mbp and 10.6 Mbp, respectively. L90 is 26, which is close to the chromosome number in FAW (31), implying nearly chromosome-sized scaffolds in this assembly. BUSCO analysis^30^ demonstrates that this assembly has the highest correctness among all published FAW genome assemblies (Table S1).

### The origin of invasion

We performed whole genome sequencing from FAW samples collected in Benin (39 individuals), India (14), Mexico (26), Florida (24), French Guiana (3), and Guadeloupe (4) using novaseq 6000 with 20X coverage on average (Fig. S1). This dataset was combined with resequencing data from populations collected in Mississippi (17) and Puerto Rico (15), which were used in our previous studies^31,32^. In addition, we had the opportunity of using resequencing data of Brazil (10), Malawi (16), and Uganda (7) from CSIRO^28^, Australia. Lastly, two individuals from China were also added to the dataset^33^. The resulting total number of individuals used in this study is 177 (99 from native populations and 78 from invasive populations). The mapping of genomes was performed against the reference assembly (Fig. S2), followed by variant calling using GATK^34^. After filtering, 27,117,672 SNPs remained (see methods for more detail). We identified the strain from a maximum likelihood phylogenetic tree reconstructed from the full sequences of mitochondrial Cytochrome C Oxidase subunit I (COX1) gene, which is the universal barcode gene and also commonly used for FAW strain identification^35^. The COX1 phylogenetic tree shows high bootstrapping confidence scores for both sfC and sfR (bootstrap supporting value > 92%) (Fig. S3), with 99 and 78 and individuals being assigned to sfC and sfR, respectively. The invasive populations have 29 and 49 sfC and sfR individuals, respectively, and native populations have 70 and 29 sfC and sfR individuals, respectively.

A principal component analysis was performed from nuclear genome sequences to identify the origin of invasive populations. The first principal component shows three groups of individuals (Fig. 1A). The first group (sfR group) consists of sfR from the Caribbean, including Florida, Guadeloupe, and French Guiana, but also one individual from Mississippi. The second group (sfC group) consists of sfC from Mexico only. The third group (hybrid group) is found between the first and second groups along the first principal component, suggesting that this group was probably generated through intraspecific hybridization between sfC and sfR. The second principal component separates the hybrid group into native (Mississippi, Puerto Rico, Brazil, and Florida) and invasive (Benin, Malawi, Uganda, India, and China) populations. This result shows that hybrids were first generated in native populations and that these hybrids further invaded the Old World. This result is in line with previous studies, indicating that the vast majority of individuals of invasive populations are hybrids^25–27^. We also observed that both native hybrid populations and invasive populations exhibit reproductive barriers between sfC and sfR from genetic differentiation (F_ST_) with the statistical significance (Fig. S4).

**Figure 1.**
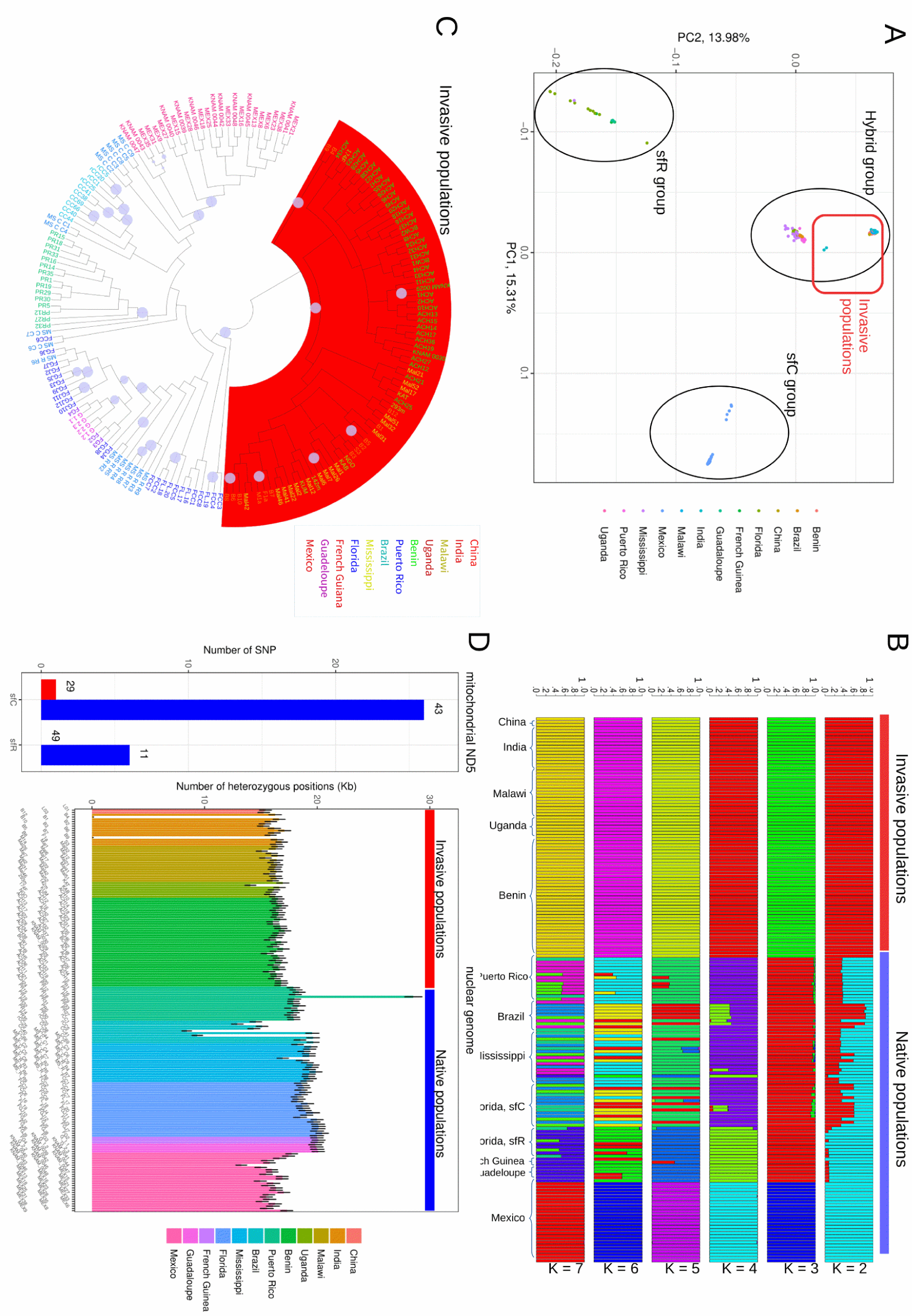
Population structure of fall armyworms. A. Principal component analysis. B. Ancestry coefficient analysis with varying K values. C. BIO-NJ phylogenetic tree was reconstructed from the allelic differentiation between a pair of individuals with 1,000 replication of bootstrapping. The circles on the branches show bootstrapping support higher than 90%. D. (left) The numbers of SNPs on the mitochondrial ND5 gene in sfC and sfR. The numbers above the bars indicate the number of sequences. (right) The number of heterozygous positions counted from positions of which genotypes are determined from all individuals. The error bars indicate 95% confidence intervals calculated from 1,000 times of bootstrapping replications in the way of resampling from 100kb windows.

The ancestry coefficient analysis^36^ shows that invasive populations have homogeneous genomic sequences in a range of K values, while native populations show the heterogeneity except for sfC in Mexico (Fig. 1B). The BIO-NJ phylogenetic tree reconstructed from whole genome sequences exhibits 100% bootstrapping supports for sfC and sfR groups (Fig. 1C), in like with the PCA results (Fig. 1A). In addition, the tree also demonstrates that all invasive individuals belong to a single clade with bootstrap support of 100%, further highlighting the homogeneity of the invasive genomic background in all the locations of collections.

### Reduction in genetic diversity during the invasion

We then compared the genetic diversity between native hybrid populations and invasive populations. We assembled whole mitochondrial genomes, and we observed that we were able to extract high-quality full-length ND5 and COX1 sequences from all 177 individuals. In ND5, the longest gene in the mitochondrial genome, sfC and sfR of the native hybrid populations have 26 and six polymorphic sites, respectively (Fig. 1D, left). However, sfC of the invasive populations has only one polymorphic site from 29 individuals (96.6% reduction), and sfR of the invasive populations has no polymorphic site from 43 individuals (100% reduction). We also compared π (nucleotide diversity) between invasive populations and native hybrid populations from whole mitochondrial genomes. The nucleotide diversity of sfC and sfR was reduced during the invasion by 78.32% (6.100 × 10^-4^ and 1.323 × 10^-4^ for native hybrid populations and invasive populations, respectively) and by 78.45% (3.156 × 10^-4^ and 6.801 × 10^-5^ for native hybrid populations and invasive populations, respectively), respectively. We identified eight and nine mitochondrial SNVs from sfC and sfR, respectively, but none of them was identified from native hybrid groups. This result implies that the observed SNVs in invasive populations were generated after the invasion, although we cannot exclude the possibility that these SNVs were derived from native hybrid populations that are not included in this study. The dramatic reduction in the mitochondrial genetic diversity, which was already shown in a previous study^37^, implies a severe genetic bottleneck during the invasion.

We further compared the number of nuclear biallelic heterozygous sites counted from each individual between native hybrid populations and invasive populations. We considered sites only if the genotype is determined from all 177 individuals to avoid potential statistical artifacts from missing data. Invasive populations have significantly lower numbers of heterozygous positions (Wilcoxon rank-sum test, *p* = 1.2 × 10^-14^), while the average difference is only 12.71% (15,854.18 and 18,162.53 for invasive populations and native hybrid populations, respectively, among 412,404bp) (Fig. 1D, right). Interestingly, two individuals from India show almost the complete depletion of heterozygosity (B4 and B9), and one individual from Puerto Rico (PR19) has particularly high heterozygosity. The dramatic difference in the reduction of genetic diversity between mitochondrial and nuclear genomes suggests that the evolutionary forces reshaping polymorphism patterns is different between these two genomes.

Multiple introductions have been suggested to contribute to an increase in the heterozygosity of invasive populations. Thus, multiple origins of FAW might explain the moderate level of heterozygosity in invasive populations. However, this explanation alone cannot explain the difference between nuclear and mitochondrial patterns shown in Fig. 1D, because it is not realistic that the admixture increased only nuclear genetic diversity (which is heterozygosity in the case of diploid nuclear genomes) while mitochondrial genetic diversity remained unchanged.

Instead, we postulate that genomic balancing selection increased nuclear heterozygosity in invasive FAW populations. In this scenario, (i) a severe bottleneck of an initially invasive population depleted heterozygosity, which caused inbreeding depression^4^ (for example, reduced egg viability, increased mortality, and reduced life span as shown in inbred monarch butterflies^38^), (ii) this population had a lagging period where the nuclear heterozygosity gradually increased through genomic balancing selection, which facilitated admixture among populations with different invasive origins, while mitochondrial genetic diversity remained low, and (iii) when the heterozygosity has sufficiently increased to generate a stable population of the initially invasive population, the FAW was able to start its large scale invasion of the Old World.

### Genomic balancing selection

To test the possibility of genomic balancing selection, we analyzed copy number variations (CNVs) to identify the origin of the invasive population with a higher resolution. As CNVs are much rarer than SNVs, we expected that CNVs have fewer noise signals from shared ancestral polymorphisms among multiple native populations to detect the invasive origin. We used CNVs only if minor allele frequency is higher than 0.2 to minimize false positives. The number of identified CNVs is 22,915. Ancestry coefficient analysis shows that, from a range of K values, invasive populations are divided into two groups (Fig. 2A). The first group includes Benin and India, and the second group includes Uganda, Malawi, and China. The first and the second groups have the same ancestry pattern from sfC in Florida (Florida-sfC) and Brazil, respectively. This result demonstrates the occurrence of multiple introductions from Florida-sfC and Brazil.

**Figure 2.**
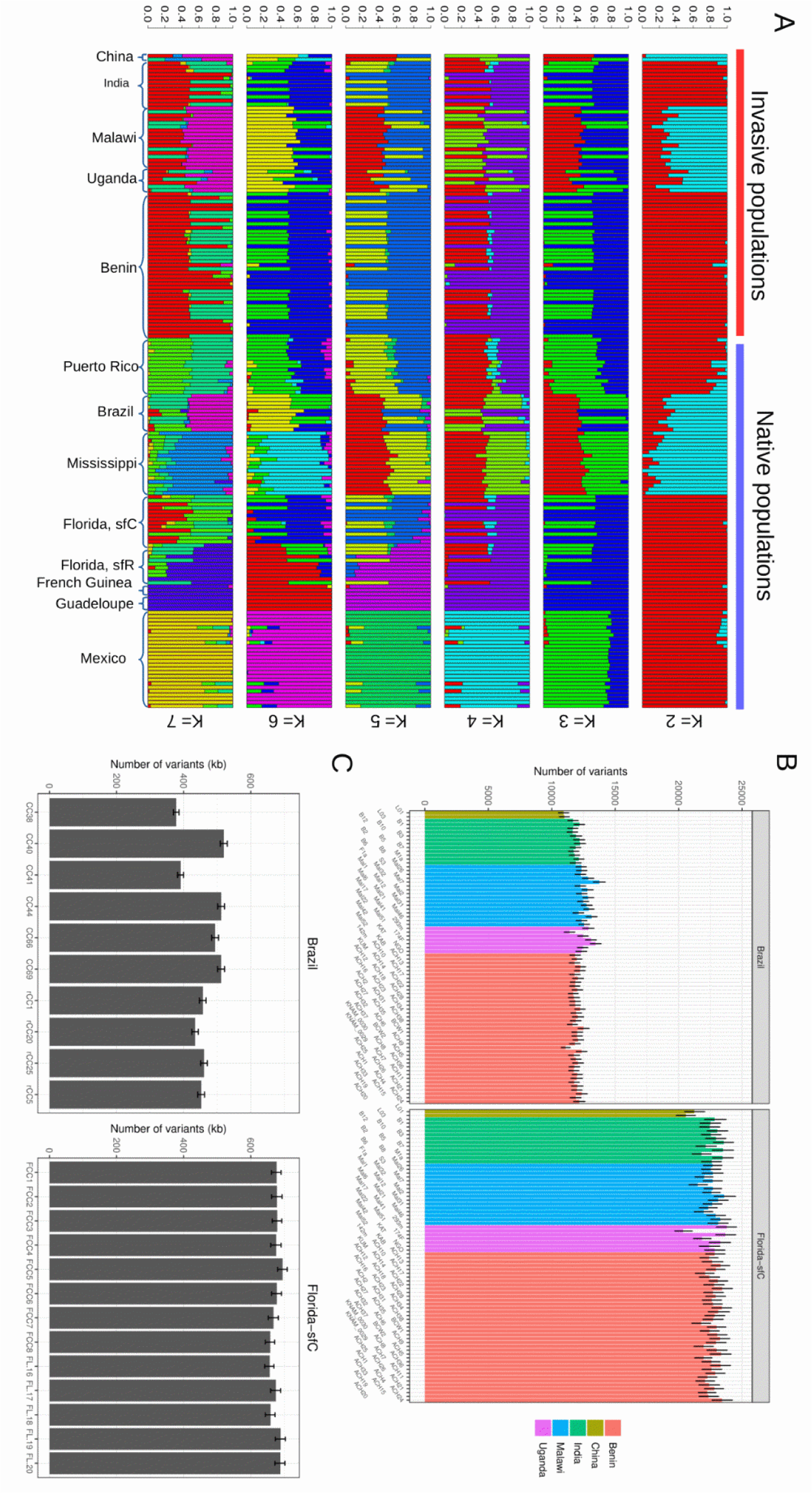
Multiple introduction of invasive fall armyworm. A. Ancestry coefficient analysis of CNV with varying K values. B. (left) The number of SNVs specifically found from the population in Brazil and absent from all the other populations, counted from each individual in the invasive populations. (right) The number of SNVs specifically found from sfC-Florida and absent from all the other populations, counted from each individual from invasive populations. C. (left) The number of SNVs in each of individuals from Brazil that are not found from sfC-Florida. (right) The number of SNVs in each of individuals from sfC-Florida that are not found from Brazil. The error bars indicate 95% confidence intervals calculated from 1,000 times of bootstrapping replication in the way of resampling from 100kb windows.

The heterogeneous distribution of Florida-sfC-specific or Brazil-specific SNV among invasive individuals was tested. We counted the numbers of SNVs that are found only from Florida-sfC or Brazil for each individual in invasive populations, and these numbers were compared between the two invasive groups (Benin-India and Malawi-Uganda-China). Fig. 2B shows a nearly uniform distribution of SNV numbers specific to Florida-sfC-specific SNV across the entire invasive populations, and the SNV numbers were not significantly different between these two groups (*p* = 0.3502; 22,746.69 and 22,493.56 for Benin-India and Malawi-Uganda-China). The Malawi-Uganda-China group has a significantly higher number of Brazil-specific SNV than the Benin-India group (*p* = 6.519 × 10^-7^; 11,934.20 and 12,484.84 for Benin-India and Malawi-Uganda-China, respectively), but with only a 4.61% difference between the two. These results show an almost uniform distribution of the numbers of Florida-sfC-specific or Brazil-specific SNVs among individuals in invasive populations, unlike what is found with CNVs.

Subsequently, we estimated to what extent the heterozygosity can be increased by admixture from SNVs that are absent in Brazil for each individual from Florida-sfC, assuming that these SNVs may increase the genetic diversity compared with a case that only Brazil is the only invading population. The numbers of these SNVs range from 656,760bp to 695,100bp (Fig. 2C). We also identified SNVs that are absent in Florida-sfC for each individual from Brazil. The numbers of these SNVs range from 378,299bp to 520,133bp. This result shows that the admixture between Florida-sfC and Brazil populations may increase the number of SNPs from 378kb to 695kb. The number of heterozygous positions in the invasive population is 1,629,133bp on average. Thus, the mixture might contribute to the heterozygous positions up to 42.67% of total invasive SNPs (695,100bp / 1,629,133bp).

Then, we tested whether genomic balancing selection increases the level of heterozygosity by mixing genes between sfC-Florida and Brazil. In the presence of balancing selection, the length of the linkage disequilibrium is decreased because balancing selection has the same effect on the linkage disequilibrium with recombination hotspot^39^. Therefore, if invasive populations experienced genomic balancing selection, then these populations are expected to have shorter linkage disequilibrium than native populations. If balancing selection does not exist, invasive populations will have longer lengths of linkage disequilibrium than native populations because of smaller effective population sizes (i.e., smaller heterozygosity as shown in Fig. 1D). To test these alternative hypotheses, we compared the decay curve of linkage disequilibrium according to the distance from one locus to another for each strain of each population. We observed that the sfC and sfR from Benin had a faster decay of linkage disequilibrium than the other invasive populations as well as sfC-Florida or Brazil populations (Fig. 3A). When the decay of linkage disequilibrium was compared across all the native and invasive populations, sfC and sfR from Benin exhibit the fastest rate of decay (Fig S5). This result shows that the invasive population in Benin has a shorter linkage disequilibrium than native populations despite the smaller effective population size. This pattern is best explained by balancing selection that increases the genomic heterozygosity level of the population in Benin. Figure 3B shows a correlation of nucleotide diversity calculated from 100kb windows between invasive and native hybrid populations. The Pearson’s correlation coefficient is very high (r = 0.992, p < 2.2 × 10^-16^), and outliers of this correlation are not observed. This pattern is in line with genomic balancing selection, rather than balancing selection affecting only a few loci. The shorter length of linkage disequilibrium in Benin is of particular interest because the FAW invasion was first reported on the Western coast of Africa, including Benin, Togo, Nigeria, and São Tomé and Príncipe^20^. Thus, we concluded that FAWs had increased heterozygosity by balancing selection in Benin (or other neighboring regions) and were able to spread eastward once their heterozygosity was sufficiently high.

**Figure 3.**
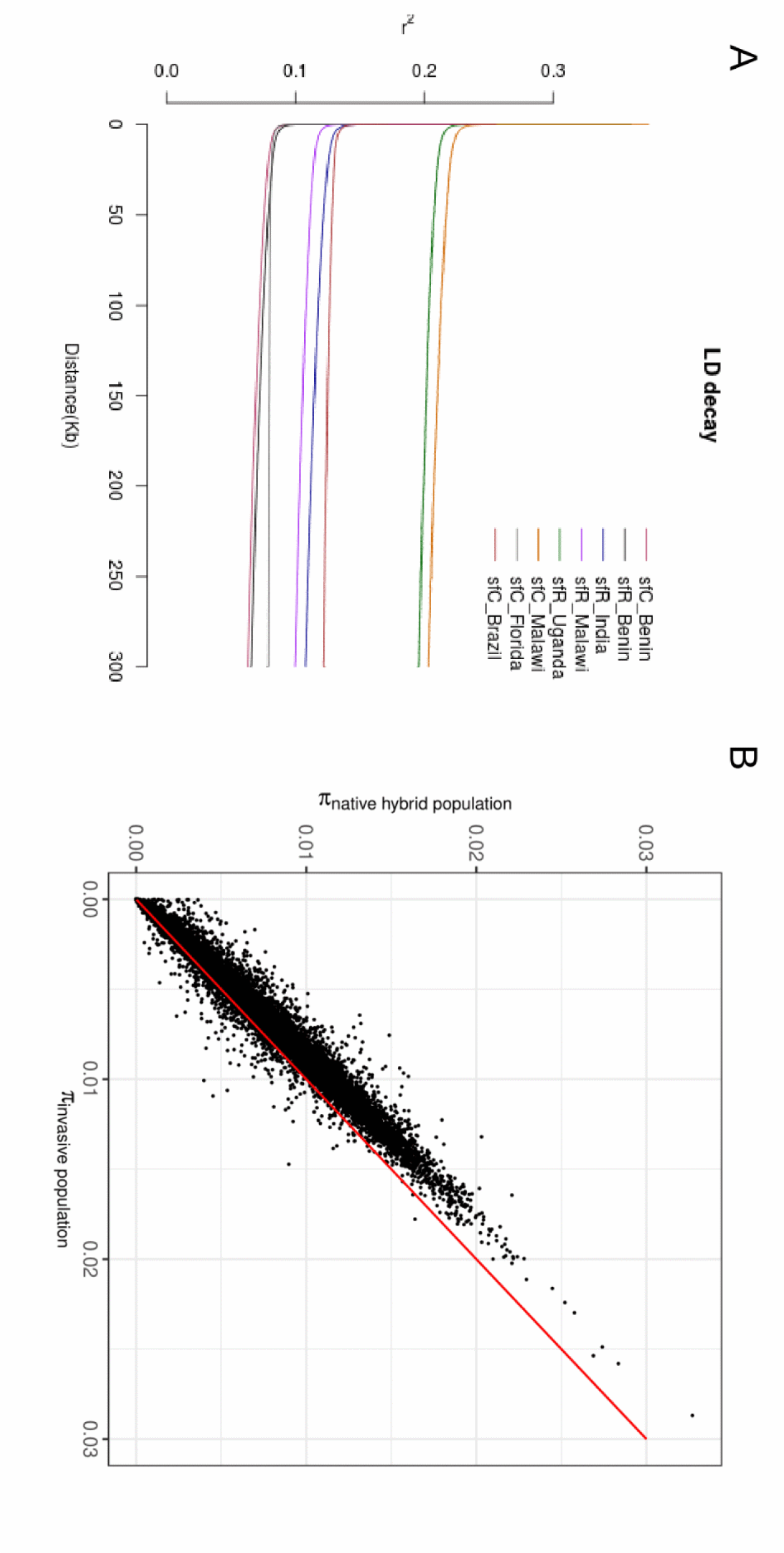
Genomic balancing selection. A. The LD decay curves calculated from each strain in each invasive population and their origins (sfC_Brazil and sfC_Florida). B. Correlation of nucleotide diversity between invasive populations and native hybrid population.

### Testing alternate hypotheses

An alternative explanation is that Florida-sfC-originated individuals co-existed with Brazil-originated individuals in Benin, while the admixture was incomplete compared with the other invasive populations. In this case, the heterogeneous genomic sequences among individuals in Benin may cause an underestimation of the length of linkage disequilibrium. We tested the heterogeneity in the population from Benin from CV (coefficient of variance) of Florida-sfC or Brazil derived variants (Fig. 2B) among invasive populations assuming that this heterogeneity among individuals increases the variance of Florida-sfC-specific or Brazil-specific SNV numbers. For the variants from Florida-sfC, CV was lowest in Benin (0.0194), followed by Malawi (0.0231), India (0.0241), and then Uganda (0.0535). For the variants from Brazil, CV was lowest in India (0.0184), followed by Benin (0.0202), Malawi (0.0332), and then Uganda (0.0527). This result shows that the population from Benin does not have a particularly high CV. Therefore, the heterogeneity of genomic sequences in Benin is not supported.

We then tested another alternative hypothesis that the level of heterozygosity in invasive populations is increased by interspecific hybridization with non-FAW species belonging to the same genus as there are several other *Spodoptera* species which are found in Africa and Asia including *S. littoralis* (Boisduval) in Africa, *S. mauritia* (Boisduval) and *S. litura* (Fabricius) in Asia, and *S. cilium* (Guenée) and *S. exigua* (Hübner) in Africa and Asia. In this case, the distribution of genetic differentiation is expected to show a bimodal distribution^40^, in which each mode represents the FAW and non-FAW species, respectively. The histogram of F_ST_ calculated from 100kb windows shows a unimodal distribution, in which 99.0% of windows have F_ST_ greater than zero (Fig. 4A). This distribution does not support inter-specific hybridization. We also tested the interspecific hybridization from the numbers of homozygous variant positions, which are expected to be increased by interspecific hybridization because, in this case, the non-FAW species have a longer phylogenetic distance from organism used to generate the reference genomes than the FAW in the native populations. In order to remove statistical artifacts, we considered positions only if genotypes are determined from all individuals. We observed that invasive populations have lower numbers of homozygous variant positions than native populations (2954.295bp and 3170.527bp in total 412,404bp for invasive and native populations, respectively; *p* = 0.005319 Wilcoxon rank-sum test) (Fig. 4B), further showing that the interspecific hybridization between *Spodoptera* species is not supported.

**Figure 4.**
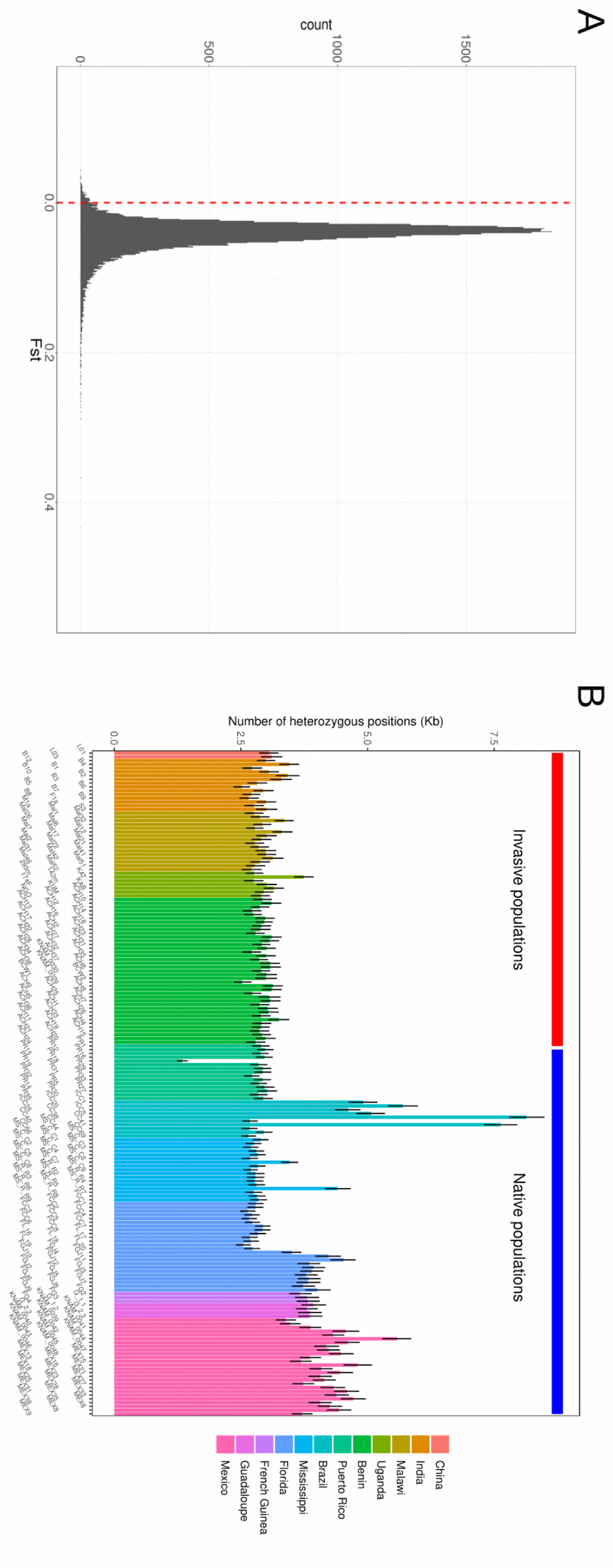
Testing interspecific hybridization. A. Histogram of F_ST_ calculated from 100kb windows between invasive populations and native hybrid groups. The red vertical bar indicates F_ST_ equals to zero. B. Homozygous variant positions were counted for each individual. The error bars indicate 95% confidence intervals calculated from 1,000 times of bootstrapping replication in the way of resampling from 100kb windows.

### Identification of adaptive evolution in the invasive population

We calculated the composite likelihood of selective sweeps^41^ from invasive populations to identify positively selected genes that may contribute to adaptation in a new environment. The median value of the composite likelihood is 0.4350, and a locus is considered to be targeted by selective sweep if the composite likelihood is higher than 100, which was arbitrarily chosen. In total, we identified seven loci on three chromosomes as potential targets of selective sweeps (Fig. 5A). As the high composite likelihood of these loci might be generated by selective sweeps not specific to invasive populations or by background selection^42^, we calculated the composite likelihood from native hybrid populations as well. Four out of the seven loci do not exhibit outliers of the composite likelihood in native hybrid populations (Fig. S6). Therefore, we considered these four loci potentially targeted by selective sweeps specific to invasive populations. These four loci contain 36 predicted protein-coding genes (Table S3), including 12 genes with unknown gene functions. We carefully underwent a manual curation of these genes to determine the function. The locus on chromosome 14 has CYP9A, which belongs to Cytochrome P450 gene family. This gene family plays a key role in detoxifying xenobiotics^43^, and CYP9A genes are overexpressed by plant allelochemicals and pesticides in FAW^44^. Therefore, positive selection on this gene might contribute to the adaptation to plants or pesticides in an invasive area. This locus also includes three copies of tubulin genes, implying that the cytoskeleton could be under positive selection as well.

**Figure 5.**
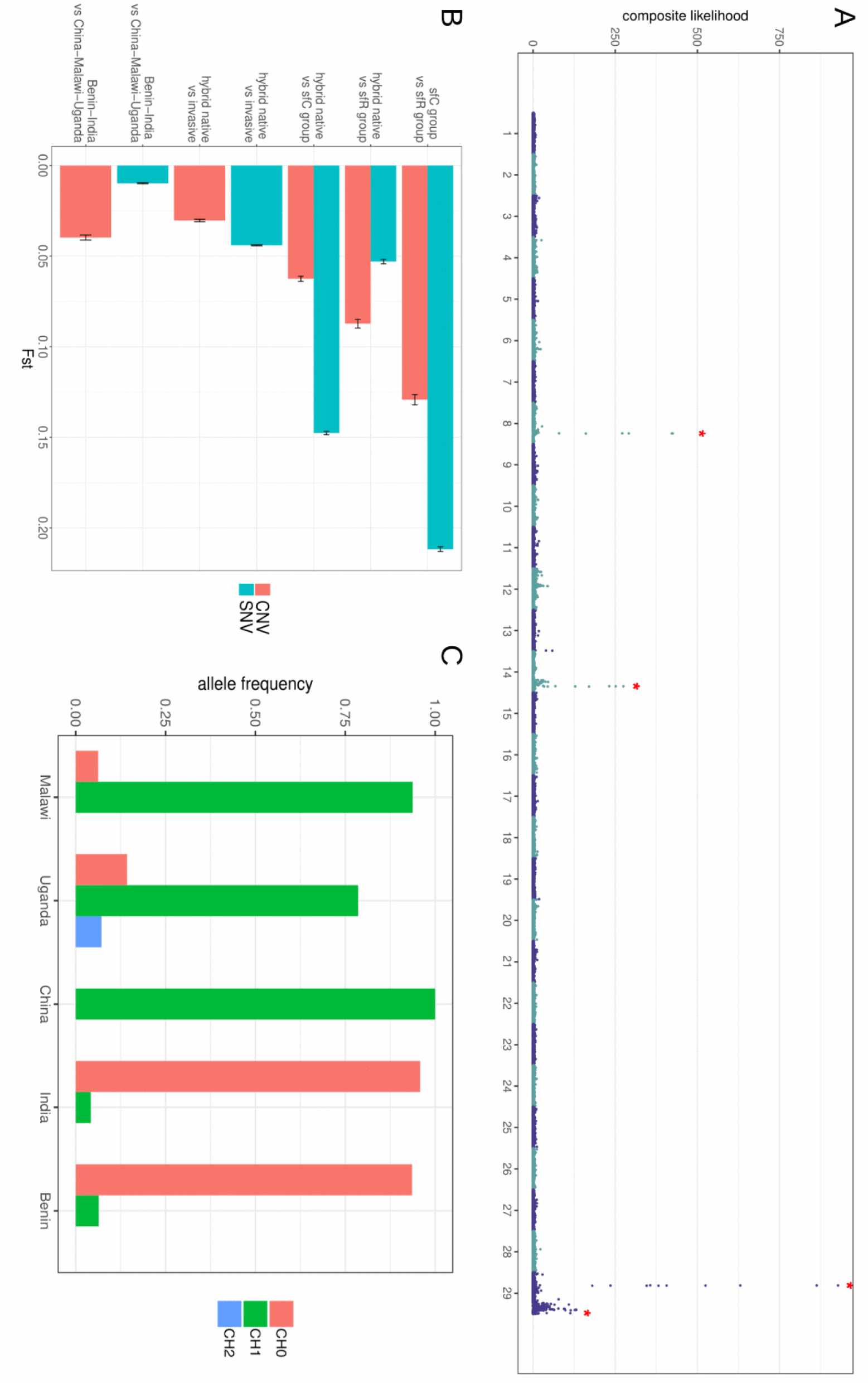
Loci under positive selection. A. The composite likelihood of being targeted by selective sweeps in invasive populations. The red asterisks indicate invasive population-specific outliers of the composite likelihood (>100), potentially targeted by selective sweeps. B. F_ST_ calculated from pairs of groups in CNV and SNV. The error bars indicate 95% confidence intervals calculated from 1,000 times of bootstrapping replication in the way of resampling from 100kb windows. C. Allele frequency of the CNV locus containing the DDSS gene. CH0, CH1, and CH2 indicate zero, one, and two copies in a haploid genome, respectively.

A locus on chromosome 29 includes a carboxylesterase gene, which may involve insecticides resistance^45^, and an ABC transporter homolog to mdr49, which protects organisms from cytotoxic compounds in *Drosophila melanogaster* Meigen^46^. Therefore, positive selection of these three genes might mitigate environmental stresses in an invasive area. This locus also includes a kunitz-type serine protease inhibitor gene, which plays a role in the digestion of plants^47^. The gene encoding odorant receptor 13, which could be important for the selection of foraging or oviposition sites^48^, is also found from this loci. Invasive populations have reduced host plant ranges compared with native populations^27,49^. One of the possible explanations of this reduction is the genetic differentiation of the serine protease inhibitor gene or the odorant receptor gene by genetic linkage to selectively targeted carboxylesterase and mdr49 genes. In this explanation, the reduction of host-plant ranges is a by-product of the process of adaptive evolution to reduce environmental stress. However, the possibility of divergent selection on the host plant should be considered as well. Interestingly, this locus includes clk, a key circadian clock gene^50^. African populations of FAWs have an earlier mating time than American populations by three hours^51^. The genetic differentiation of clk could also be caused by genetic linkage to positively selected environmental stress genes or host-plant genes, while divergent selection on the circadian clock is also possible.

CNV exhibits two groups in the invasive population (Fig. 2A), unlike SNV. The first group includes Benin and India, and the second group includes China, Malawi, and Uganda. We tested the presence of positive selection by CNV that is specific to one or both groups in invasive populations. F_ST_ calculated from CNV between Benin-India and China-Malawi-Uganda is 0.0397 (Fig. 5B). F_ST_ calculated from SNV between these two groups is 0.00973, which represents only 24.5% of F_ST_ from F_ST_ from CNV (0.00973/0.0397). In order to test if CNV having much higher F_ST_ than SNV is a general phenomenon, we also calculated F_ST_ between pairs among native hybrid populations, sfC group, and sfR group. The ratio of F_ST_ between these pairs from SNV to CNV ranges from 0.607 to 2.36 (Fig. 5C), which is higher than the ratio of F_ST_ between Benin-India and China-Malawi-Uganda (0.245). Thus, we concluded that the F_ST_ calculated from CNV between Benin-India and China-Malawi-Uganda could be affected by positive selection on CNV. In total, six loci with CNV have almost complete genetic differentiation between the two groups (F_ST_ > 0.8).

We identified only one gene, Decaprenyl-diphosphate synthase subunit 2 (DDSS2), from these loci. Most individuals in the China-Malawi-Uganda group have this gene as single-copy, while the Benin-India group lacks this gene in most individuals. In FAW, the DDSS gene is down-regulated by bat-induced stress^52^, and a region near Benin exhibits a hotspot for bat-species diversity^53^. Thus, the CNV of DDSS gene could possibly be a consequence of adaptation to local bat communities in West Africa (or India). More ecological studies are required to test the differential stress from predators across multiple invasive populations.

In this study, we showed that the restoration of the level of heterozygosity by genomic balancing selection is key to invasive success in FAW and that it likely enables its rapid global invasion of the Old World. We do not argue that invasive FAW in Western Africa obtained a new trait by adaptive evolution that increased invasiveness (e.g., Bridgehead Effects^54,55^). FAWs in native populations exhibit high migratory behavior, and invasive populations have probably equally high mobility as native populations. Instead, we argue here that the generation of a stable population in West Africa by genomic balancing selection played a key role in invasive success in FAW.

In addition, we do not argue that West Africa is the only initially invaded area. It is possible that the initial introduction of FAW might occur elsewhere in the Eastern Hemisphere^28^, while invasive FAW remained undetected due to their small population size. We argue here that genomic balancing selection is one of the causal evolutionary forces responsible for explosive population growth in West Africa by facilitating admixtures and that this population migrated eastward, as shown from the chronological order of detection of invasive FAW. If populations of FAW existed in the Eastern Hemisphere before the first detection in West Africa, potential gene flow among invasive populations could explain the different patterns of ancestry coefficients between CNV (Fig. 2A) and SNV (Fig. 1B) among invasive populations.

The majority of reported cases show that the reduction in heterozygosity is mild (e.g., < 20%) in a wide range of taxa^1^. Therefore, it could be postulated that balancing selection may play a key role in the invasive success of a large range of organisms. Future studies should involve population genomics analysis in other invasive taxa to test this possibility. This study also highlights the importance of rapid and vigorous pest control during the early phase of the invasion, as emphasized by many researchers, before heterozygosity is sufficiently increased to generate a stable population by genomic balancing selection. For an early eradication, early monitoring of pest species is mandatory, and a small number of individuals should not be overlooked, like the case of the Asian hornet (*Vespa velutina* Lepeletier) that started from a small invasive population which then went on to rapidly colonize large areas of Western Europe^56^.

### Methods

### Genome assembly

We performed the mapping of Illumina reads (~80X)^57^ against an assembly, which was generated from 30X PacBio Reads in our previous study^32^, using SMALT^58^, and potential errors in the assemblies were identified using reapr^59^. If an error is found over a gap, the scaffold was broken into two using the same software to remove potential structural errors in the assembly. The broken assemblies were concatenated using SALSA2^60^ or 3D-DNA^61^, followed by gap filling with the 80X Illumina reads using SOAP-denovo2 Gap-Closer^62^ and with the PacBio reads using LR_GapCloser v1.1^63^. We observed that 3D-DNA generated a slightly more correct assembly than SALSA2 from BUSCO analysis (Table S1). Thus, the assembly from 3D-DNA was used in this study. Gene annotation was transferred from the previously generated assemblies to current assembly using RATT^64^.

### Resequencing Data

FAW larvae were collected from Wagou and Gando Villages in Benin (2017), from Citra and Jacksonville in Florida (2015), from Texcoco in Mexico (2009), from French Guiana (1992), and from Petit-Bourg and Port-Louis in Guadeloupe (2013). We obtained gDNA from India, which was used by Sharanabasappa et al^65^. Genomic DNA was extracted using the Wizard Genomic DNA kit or the Qiagen Dneasy blood and tissues kit. Libraries for whole genome resequencing were constructed from 1.0μg DNA per sample using NEBNext DNA Library Prep Kit. Novaseq 6000 g DNA per sample using NEBNext DNA Library Prep Kit. Novaseq 6000 with ~20X coverage was used to perform whole genome resequencing with 150bp paired-end and 300bp insert length. Then, we combined the resequencing data from Puerto Rico and Mississippi, which were generated for our previous studies (Hiseq 2500, Hiseq 4000, and Novaseq 6000)^31,32^, as well as the resequencing data of Brazil, Malawi, and Uganda from CSIRO (Novaseq 6000, 150bp paired-end sequencing)^28^. Lastly, resequencing data from China^33^ was also combined with the dataset. Adapter sequences were removed using adapterremoval^66^. Then, we performed mapping of reads against the reference genome using bowtie2^67^. Then, we performed a variant calling using GATK^34^. Filtering was performed if QD is lower than 2.0, or FS is higher than 60.0, or MQ is lower than 40.0, or MQRankSum is lower than -12.5, or ReadPosRankSum is lower than -8.0. CNVs were identified using CNVCaller^68^. We discarded all CNVs unless minor allele frequency is higher than 0.2 to reduce false positives.

### Phylogenetic analysis

To identify strains, we mapped the Illumina reads against mitochondrial genomes (NCBI: KM362176) using bowtie2^67^, followed by extracting mitochondrial reads using samtools^69^. Mitochondrial genomes were assembled using MitoZ^70^, and COX1 sequences were identified. These COX1 sequences were aligned together with a COX1 sequence from a specimen of another *Spodoptera* species, *S. exigua* (NCBI ID, JX316220), using MUSCLE ^71^, and a maximum likelihood phylogenetic tree was reconstructed using PhyML^72^. The phylogenetic tree was visualized using iTOL ^73^.

We calculated the nuclear genetic distance between each pair of individuals from the difference in allele frequency at biallelic sites in which genotypes are determined from all individuals using VCFphylo (https://github.com/kiwoong-nam/VCFPhylo). Transversional variants were weighted to two. Then, a bootstrapping distance matrix was generated with 1,000 replications, and we generated BIO-NJ trees for each matrix using FastME^74^. Then, a consensus tree was made using consense in Phylip package^75^, and the tree was visualized using iTOL^73^.

### Population genomics analysis

The principal component analysis was performed using plink^76^. We used admixture^36^ for the ancestry coefficient analysis. Weir and Cockerham’s F_ST_^77^ was calculated using VCFtools^78^. Potential targets of selective sweeps were identified using SweeD^41^. The number of the grid is 1,000 per chromosome. If a locus has the composite likelihood of selective sweeps higher than 100, we considered that this locus was targeted by a selective sweep. The decay curves of linkage disequilibrium were generated using PopLDdecay^79^. To identify mitochondrial SNVs, a mitochondrial VCF was generated from the bam files, which was made to identify strains (see above), using GATK^34^.

## Acknowledgements

This work (ID 1702-018, given to KN) was publicly funded through ANR (the French National Research Agency) under the "Investissements d’avenir" programme with the reference ANR-10-LABX-001-01 Labex Agro and coordinated by Agropolis Fondation under the frame of I-SITE MUSE (ANR-16-IDEX-0006). In addition, a grant from the department of Santé des Plantes et Environnement at Institut national de la recherche agronomique for KN (adaptivesv). This work was also financially supported by EUPHRESCO (FAW-spedcom, given to Anne-Nathalie Volkoff) and by CSIRO Health & biosecurity (given to WTT, TW, and KG). SY was supported by a CIRAD-INRAE PhD fellowship.

## Author Contributions

FL generated reference genome assembly. WTT, MF, SD, RA, CMK, RLMJ, CAB, PS, TB, AD, TW, KG, and NN provided samples for whole genome resequencing. EF, ANC, SG, and GJK prepared samples. EF performed variant calling. SY and KN performed analysis. NN and EA performed gene annotation. SY and KN wrote manuscript. KN involved in planning and supervised the work.

## Additional Information

The raw reads of these samples are available from NCBI SRA (PRJNA639296 for samples from Florida and PRJNA639295 for the rest of the samples). The reference genome assembly used in this study is available at BIPAA (https://bipaa.genouest.org/sp/spodoptera_frugiperda). Supplementary Information is available for this paper. We declare a full code availability upon request.

