## Supplementary material for "Genomic balancing selection is key to the invasive success of the fall armyworm": Table S1 - Table S3, Fig S1 - Fig S6

Table S1. Summary statistics of genome assemblies showing contiguity and correctness of the assemblies (BUSCO) generated in this study and other published assemblies.

| | Statistics | Current assembly | Nam et al.[22] | Nam et al.[23] | Gouin et al[49] | | Liu et al.[24] | |
|---|---|---|---|---|---|---|---|---|
| | | | | | sfC | sfR | Male | Female |
| Contiguity | Assembly size | 385,049,154 | 384,455,365 | 379,902,278 | 437,873,304 | 371,020,040 | 543,659,128 | 531,931,622 |
| | number of sequences | 495 | 125 | 1,054 | 41,577 | 29,127 | 21,840 | 27,258 |
| | N50 (bp) | 13,567,047 | 13,151,234 | 1,129,192 | 52,781 | 28,526 | 14,162,803 | 13,967,093 |
| | L50 | 13 | 13 | 91 | 1,616 | 3,761 | 16 | 17 |
| | N90 (bp) | 10,609,592 | 8,473,354 | 165,330 | 3,545 | 6,422 | 6,440 | 5,122 |
| | L90 | 26 | 27 | 421 | 18,789 | 13,881 | 3,030 | 5,612 |
| BUSCO | Complete | 1,601 | 1,601 | 1,616 | 1,461 | 1,551 | 1,576 | 1,577 |
| | Complete and single-copy | 1,577 | 1,573 | 1,573 | 1,276 | 1,518 | 1,442 | 1,480 |
| | Complete and duplicated | 25 | 28 | 43 | 185 | 33 | 134 | 97 |
| | Fragmented | 19 | 20 | 11 | 127 | 69 | 45 | 48 |
| | Missing | 37 | 37 | 31 | 70 | 38 | 37 | 33 |
| | Total | 1,658 | 1,658 | 1,658 | 1,658 | 1,658 | 1,658 | 1,658 |

Table S2. Summary statistics of genome assemblies showing contiguity and correctness of the assemblies generated using SALSA2 and 3D-DNA.

| | | | |
|---|---|---|---|
| Contiguity | Number of scaffolds | 390 | 495 |
| | Size | 384Mb | 385Mb |
| | N50 | 22.8Mb | 13.6Mb |
| | L50 | 6 | 13 |
| | N90 | 0.8MB | 10.6Mb |
| | L90 | 30 | 26 |
| Correctness (BUSCO) | Complete | 1,601 | 1,601 |
| | Complete and single-copy | 1,570 | 1,577 |
| | Complete and duplicated | 31 | 25 |
| | Fragmented | 22 | 19 |
| | Missing | 35 | 37 |

Table S3. The list of genes on the potential targets of invasive specific selective sweeps

| Chromosome | start | end | ID | gene name |
|---|---|---|---|---|
| 8 | 8,554,974 | 8,560,903 | SFRUCORN0000021653 | Protein of unknown function |
| 8 | 8,590,850 | 8,597,741 | SFRUCORN0000021660 | Protein of unknown function |
| 8 | 8,563,080 | 8,563,500 | SFRUCORN0000021654 | Protein of unknown function |
| 8 | 8,588,488 | 8,588,871 | SFRUCORN0000021658 | Protein of unknown function |
| 8 | 8,571,015 | 8,595,815 | SFRUCORN0000021655 | Atrial natriuretic peptide receptor 1 |
| 14 | 12,275,316 | 12,282,581 | SFRUCORN0000025984 | Cytochrome p450 CYP9A75 |
| 14 | 12,386,949 | 12,388,669 | SFRUCORN0000025990 | Tubulin beta-3 chain |
| 14 | 12,389,894 | 12,391,590 | SFRUCORN0000025991 | Tubulin beta-3 chain |
| 14 | 12,395,694 | 12,402,075 | SFRUCORN0000025993 | alpha-1,2-mannosyltransferase ALG9 |
| 14 | 12,405,090 | 12,407,146 | SFRUCORN0000025995 | Pre-rRNA-processing protein TSR2 |
| 14 | 12,391,848 | 12,393,676 | SFRUCORN0000025992 | Tubulin beta-3 chain |
| 14 | 12,320,781 | 12,331,540 | SFRUCORN0000025986 | Protein of unknown function |
| 14 | 12,371,977 | 12,379,028 | SFRUCORN0000025988 | Protein of unknown function |
| 14 | 12,404,482 | 12,405,037 | SFRUCORN0000025994 | Protein of unknown function |
| 14 | 12,407,580 | 12,429,248 | SFRUCORN0000025996 | DNA topoisomerase 2-binding protein 1 |
| 29 | 6,570,648 | 6,570,917 | SFRUCORN0000001059 | Probable Transposon |
| 29 | 6,646,496 | 6,651,061 | SFRUCORN0000001051 | Protein of unknown function |
| 29 | 6,596,394 | 6,616,494 | SFRUCORN0000001055 | peptide transporter |
| 29 | 6,654,052 | 6,658,901 | SFRUCORN0000001050 | Protein of unknown function |
| 29 | 6,582,434 | 6,589,323 | SFRUCORN0000001057 | Fer 2 homolog |
| 29 | 6,644,505 | 6,645,033 | SFRUCORN0000001052 | Protein of unknown function |
| 29 | 6,598,209 | 6,599,921 | SFRUCORN0000001056 | carboxylesterase 022a (cxe022a) |
| 29 | 6,573,145 | 6,577,083 | SFRUCORN0000001058 | palmytoyltransferase |
| 29 | 6,693,319 | 6,704,962 | SFRUCORN0000001044 | centrosomal protein |
| 29 | 6,660,376 | 6,665,474 | SFRUCORN0000001048 | odorant receptor 13 |
| 29 | 6,692,263 | 6,692,466 | SFRUCORN0000001045 | Protein of unknown function |
| 29 | 6,671,207 | 6,673,259 | SFRUCORN0000001047 | SPARC-related modular calcium-binding protein |
| 29 | 6,683,531 | 6,691,698 | SFRUCORN0000001046 | SPARC-related modular calcium-binding protein |
| 29 | 6,710,652 | 6,729,471 | SFRUCORN0000001043 | ubiquitin carboxyl-terminal hydrolase |
| 29 | 6,654,220 | 6,659,767 | SFRUCORN0000001049 | Kunitz-type serine protease inhibitor |
| 29 | 6,530,682 | 6,542,329 | SFRUCORN0000001062 | Clock (CLK) |
| 29 | 6,548,052 | 6,549,563 | SFRUCORN0000001061 | Protein of unknown function |
| 29 | 6,554,517 | 6,567,412 | SFRUCORN0000001060 | Uncharacterized, RING domain protein family |
| 29 | 6,513,073 | 6,526,354 | SFRUCORN0000001063 | ABC transporter; multidrug resistance protein homolog 49-like |
| 29 | 20,629,938 | 20,637,386 | SFRUCORN0000010362 | zinc finger and BTB domain-containing protein |
| 29 | 20,638,804 | 20,639,796 | SFRUCORN0000010363 | UDP sugar transporter, probable UST74c |

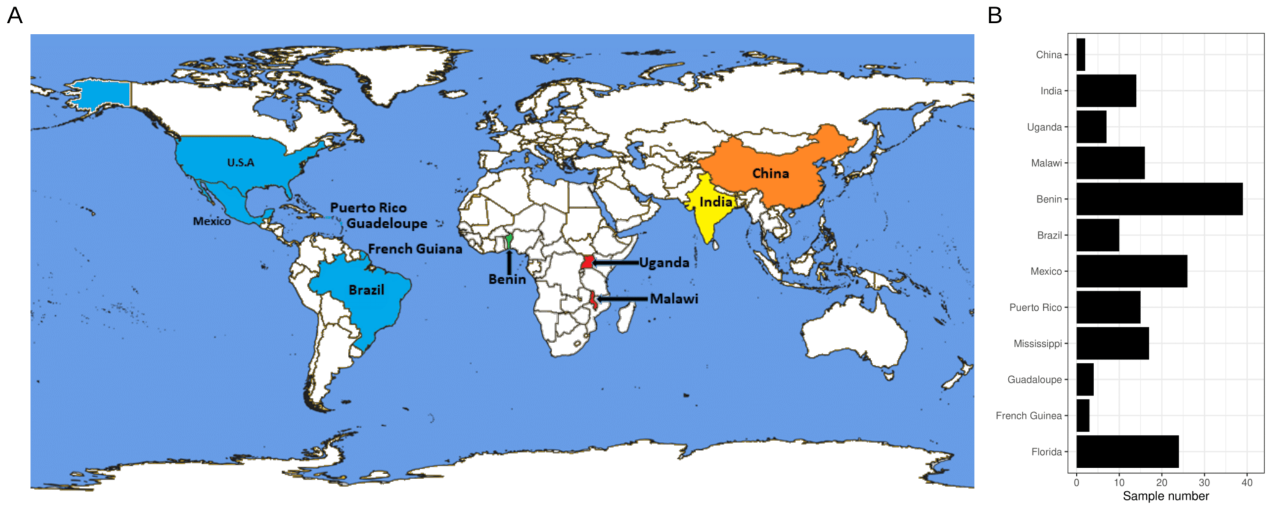


Figure S1. A. Map indicating the countries from which samples of *Spodoptera frugiperda* were sequenced. The blue color indicates the native habitat of *S. frugiperda.* The green, red, yellow, and orange colors indicate that the invasion was reported in 2016, 2017, 2018, and 2019 respectively. B. The numbers of samples used in this study.

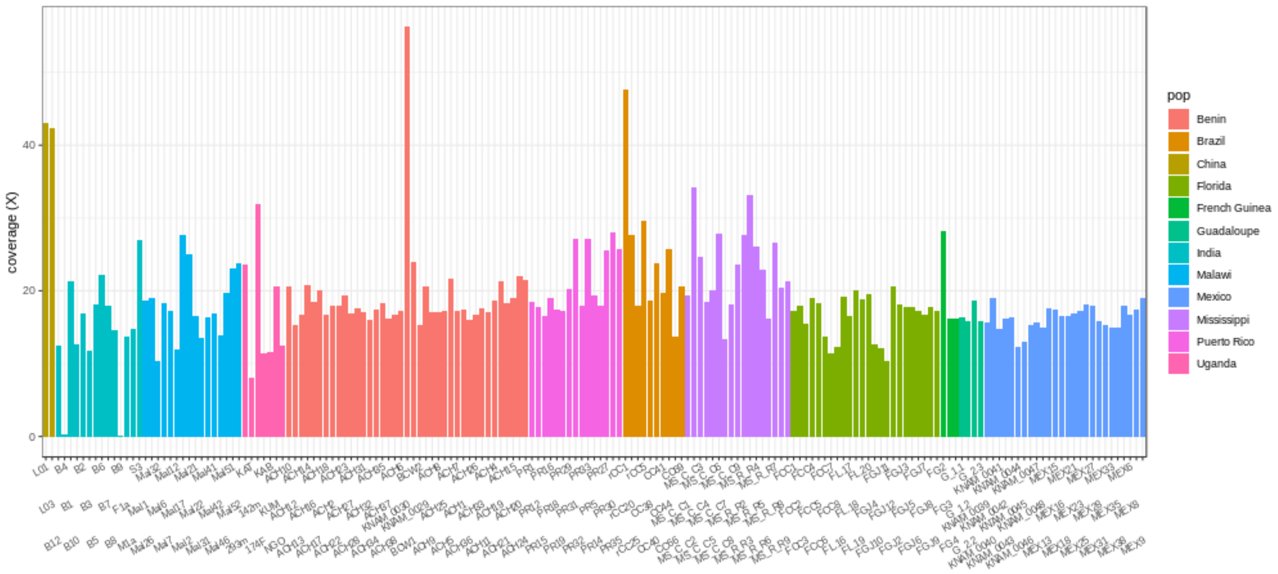


Figure S2. Read depth of each sequenced individual.

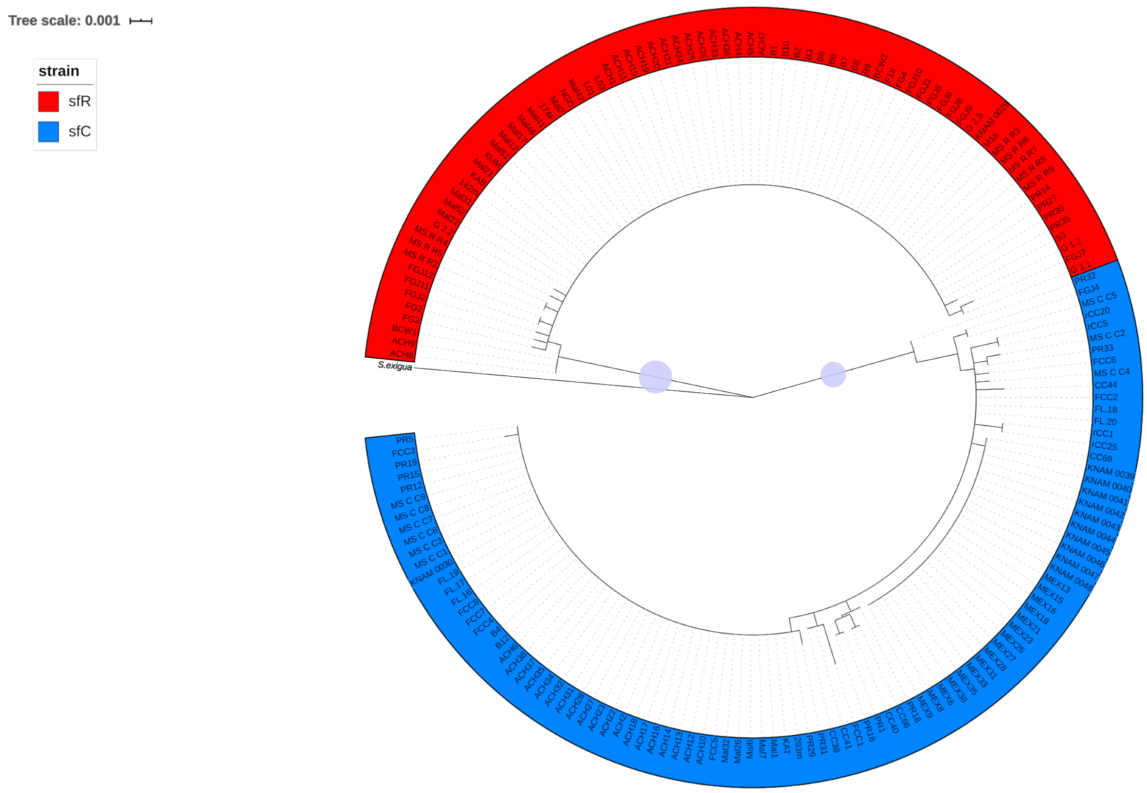


Figure S3. A maximum likelihood phylogenetic tree reconstructed from full mength mitochondrial COX1 (1,536bp). The red and blue clades indicate sfR and sfC, respectively. The circles on the branches show bootstrapping support higher than 90%. The length of the branch to *S. exigua* was reduced to 1/3 for the visualization purpose.

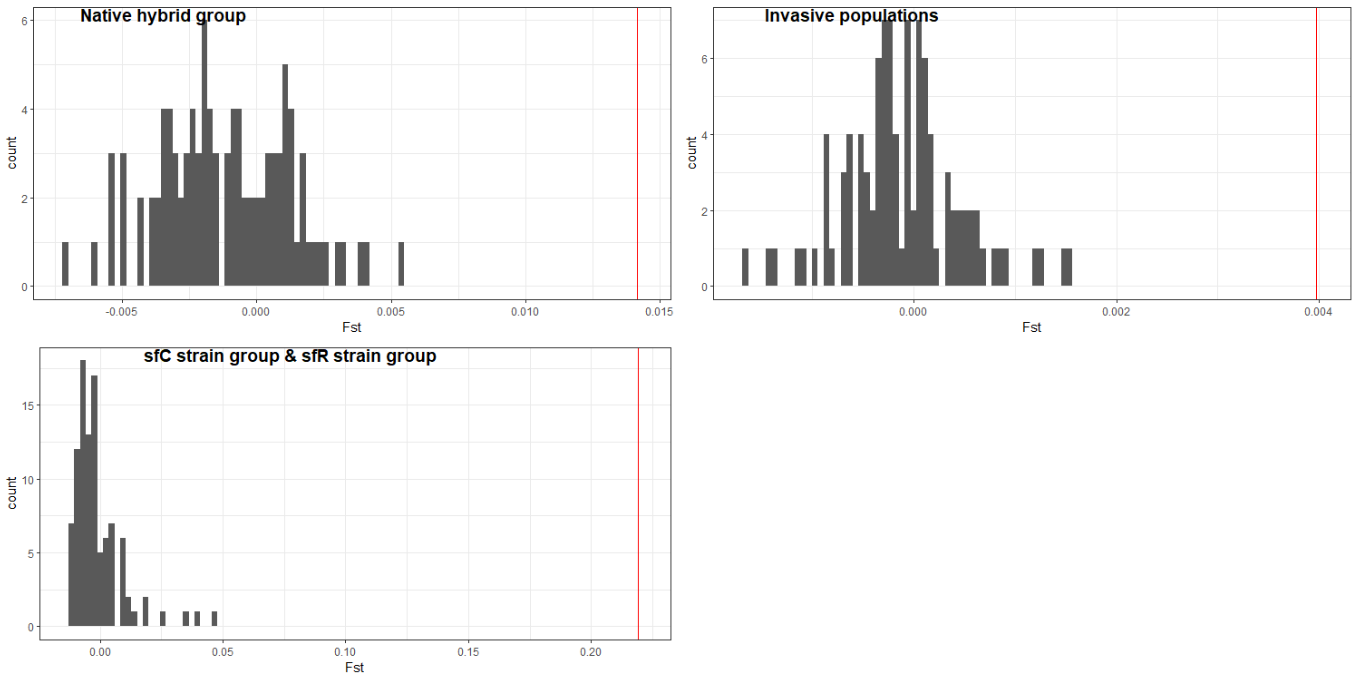


Figure S4. Significant genetic differentiation between sfC and sfR. The histograms show $F_{ST}$ calculated from randomly generated two groups in the native hybrid group, in invasive populations, and in sfC group (Mexico) and sfR group (sfR from the Caribbean). The red vertical bars indicate $F_{ST}$ calculated between sfC and sfR.

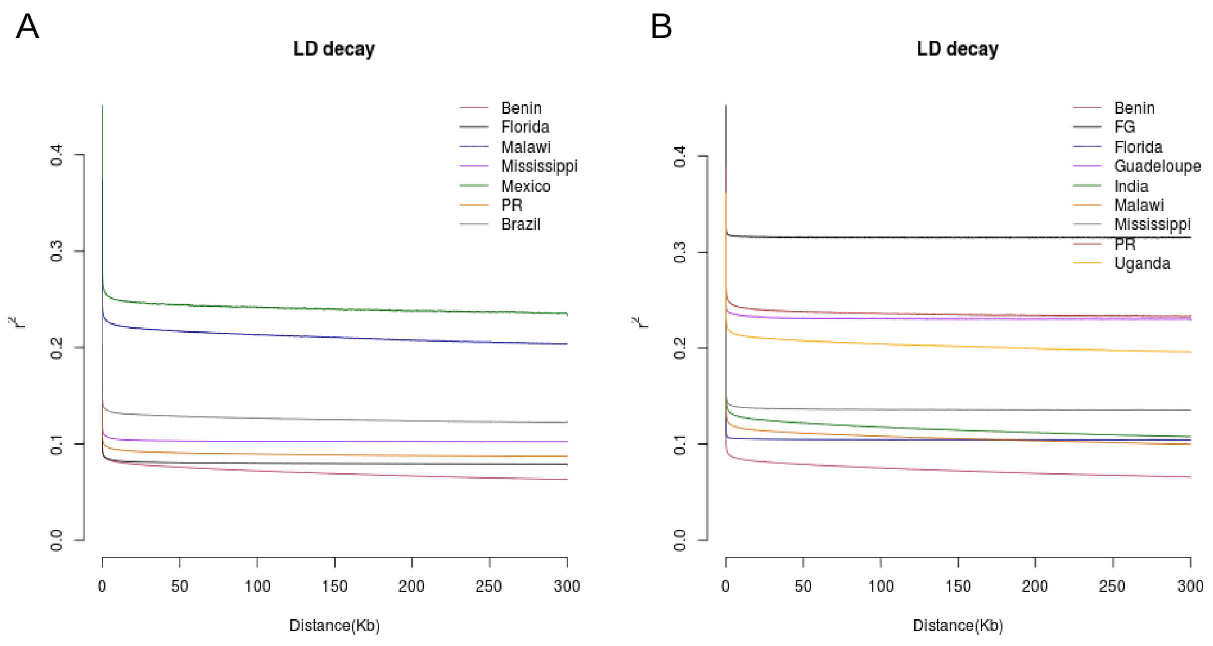


Figure 5. The LD decay curve of each geographic population in (A) sfC and (B) sfR. In both cases, the populations from Benin have the fastest decay of linkage disequilibrium among all populations.

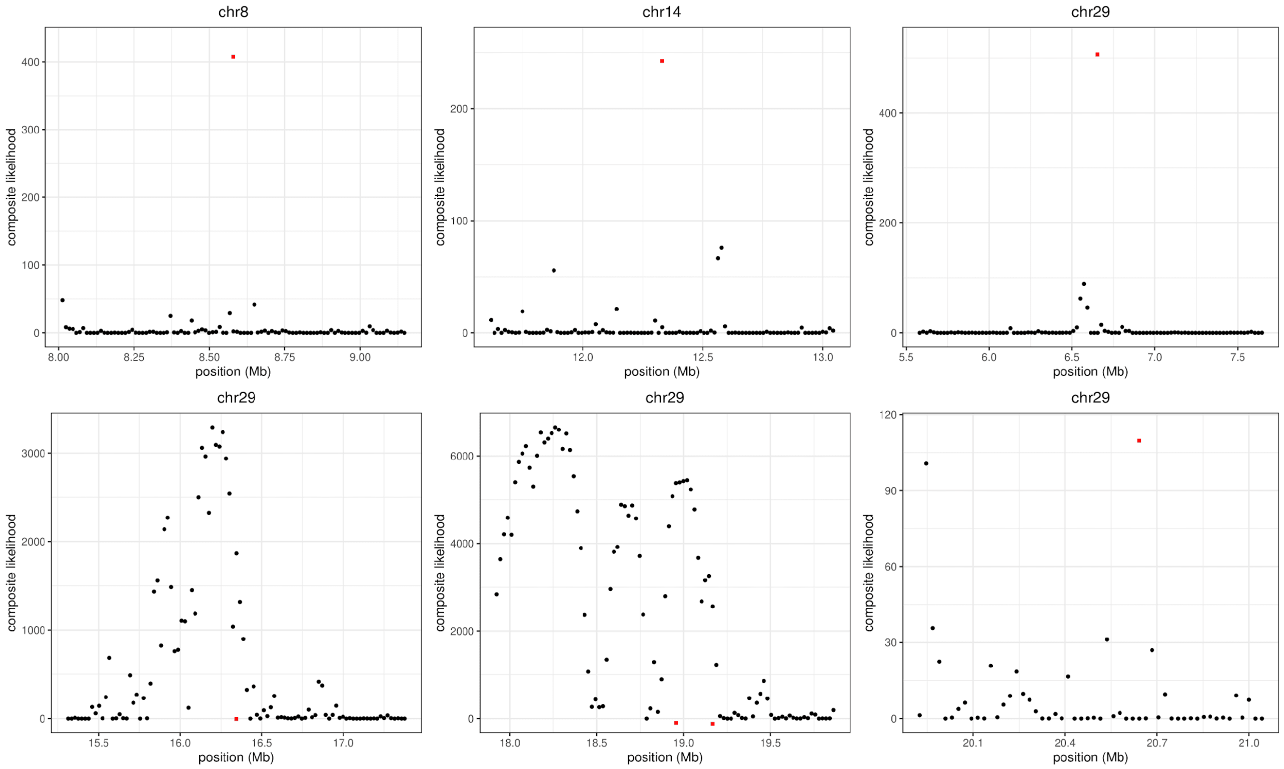


Figure S6. The distribution of composite likelihoods calculated from native hybrid populations at the identified seven outliers in invasive populations. The red dots indicate the composite likelihood of invasive populations in the middle of outliers. Native hybrid populations do not have outliers of the composite likelihoods at the first three and the last outliers. Therefore, we considered that these four loci are potential targets of selective sweeps.
